# Connectivity Problems on Heterogeneous Graphs

**DOI:** 10.1101/300012

**Authors:** Jimmy Wu, Alex Khodaverdian, Benjamin Weitz, Nir Yosef

## Abstract

**Background:** Network connectivity problems are abundant in computational biology research, where graphs are used to represent a range of phenomena: from physical interactions between molecules to more abstract relationships such as gene co-expression. One common challenge in studying biological networks is the need to extract meaningful, small subgraphs out of large databases of potential interactions. A useful abstraction for this task turned out to be the Steiner network problems: given a reference “database” graph, find a parsimonious subgraph that satisfies a given set of connectivity demands. While this formulation proved useful in a number of instances, the next challenge is to account for the fact that the reference graph may not be static. This can happen for instance, when studying protein measurements in single cells or at different time points, whereby different subsets of conditions can have different protein milieu.

**Results and Discussion:** We introduce the *condition* Steiner network problem in which we concomitantly consider a set of distinct biological conditions. Each condition is associated with a set of connectivity demands, as well as a set of edges that are assumed to be present in that condition. The goal of this problem is to find a minimal subgraph that satisfies all the demands through paths that are present in the respective condition. We show that introducing multiple conditions as an additional factor makes this problem much harder to approximate. Specifically, we prove that for *C* conditions, this new problem is NP-hard to approximate to a factor of *C* – *ϵ*, for every *C* ≥ 2 and *ϵ* > 0, and that this bound is tight. Moving beyond the worst case, we explore a special set of instances where the reference graph grows *monotonically* between conditions, and show that this problem admits substantially improved approximation algorithms. We also developed an integer linear programming solver for the general problem and demonstrate its ability to reach optimality with instances from the human protein interaction network.

**Conclusion:** Our results demonstrate that in contrast to most connectivity problems studied in computational biology, accounting for multiplicity of biological conditions adds considerable complexity, which we propose to address with a new solver. Importantly, our results extend to several network connectivity problems that are commonly used in computational biology, such as Prize-Collecting Steiner Tree, and provide insight into the theoretical guarantees for their applications in a multiple condition setting.

**Availability:** Our solver for the general *condition* Steiner network problem is available at https://github.com/YosefLab/condition_connectivity_problems

## Background

In molecular biology applications, networks are routinely defined over a wide range of basic entities such as proteins, genes, metabolites, or drugs, which serve as nodes. The edges in these networks can have different meanings, depending on the particular context. For instance, in protein-protein interaction (PPI) networks, edges represent physical contact between proteins, either within stable multi-subunit complexes or through transient causal interactions (i.e., an edge (*x*,*y*) means that protein *x* can cause a change to the molecular structure of protein *y* and thereby alter its activity). The body of knowledge encapsulated within the human PPI network (tens of thousands of nodes and hundreds of thousands of edges in current databases, curated from thousands of studies [1]) is routinely used by computational biologists to generate hypotheses of how various signals are transduced in eukaryotic cells [2, 3, 4, 5, 6]. The basic premise is that a process that starts with a change to the activity of protein *u* and ends with the activity of protein *v* must be propagated through a chain of interactions between *u* and *v*. The natural extension regards a process with a certain collection of protein pairs {(*u*_1_, *v*_1_),…, (*u_k_*, *v_k_*)}, where we are looking for a chain of interactions between each *u_i_* and *v_i_*. In another set of applications, the notion of directionality is not directly assumed and instead, one is looking for a parsimonious subgraph that connects together a set *S* of proteins that are postulated to be active [7, 8].

In most applications, the identity of the so called terminal nodes (i.e., (*u_i_*, *v_i_*) pairs or the set *S*) is assumed to be known (or inferred from experimental data such as ChIP-seq [7, 8, 5]), while the identity of the intermediate nodes and interactions is unknown. The goal therefore becomes to complete the gap and find a probable subgraph of the PPI network that simultaneously satisfies all the connectivity demands, thereby explaining the overall biological activity. Since the edges in the PPI network can be assigned a probability value (reflecting the credibility of their experimental evidence), by taking the negative log of these values as edge weights, the task becomes minimizing the total edge weight, leading to an instance of the Steiner Network problem. We have previously used this approach to study the propagation of a stabilizing signal in proinflammatory T cells, leading to the identification of a new molecular pathway (represented by a sub-graph of the PPI network) that is critical for mounting an auto-immune response, as validated experimentally by perturbation assays and disease models in mice [5]. Tuncbag et al. [8] have utilized the undirected approach using the Prize-Collecting Steiner Tree model, where the input is a network *G* along with a penalty function, *p*(*v*) for each protein (node) in the network (based on their importance; e.g., fold-change across conditions). The goal in this case is to find a probable subtree which contains the majority of the high cost proteins in *G*, while accounting for penalties paid by both edge usage and missing proteins, in order to capture the biological activity represented in such a network [7, 8].

While these studies contributed to our understanding of signal transduction pathways in living cells, they do not account for a critical aspect of the underlying biological complexity. In reality, proteins (nodes) can become activated or inactivated at different conditions, thereby giving rise to a different set of potential PPIs that might take place [9]. Here, the term *condition* can refer to different points in time [10], different treatments [11], or, more recently, different cells [12]. Indeed, advances in experimental proteomics provide a way to estimate these changes at high throughput, e.g., measuring phosphorylation levels or overall protein abundance, proteome-wide for a limited number of samples [11]. A complementary line work provides a way to evaluate the abundance of smaller numbers of proteins (typically dozens of them) in hundreds of thousands of single cells [12].

The next challenge is therefore to study connectivity problems that take into account not only the endpoints of each demand, but also the condition in which these demands should be satisfied. This added complication was tackled by Mazza et al. [13], who introduced the “Minimum *k*-Labeling (MKL)” problem. In this setting, each connectivity demand comes with a label, which represents a certain experimental condition or time point. The task is to label edges in the PPI network so as to satisfy each demand using its respective label, while minimizing the number of edges in the resulting sub-graph and the number of labels used to annotate these edges. While MKL was an important first step, namely introducing the notion of different demands for each condition, the more difficult challenge still remains that of considering variability in the reference graph, namely different sets of proteins that may be active and available for use in each condition.

## Summary of Main Contributions

To address this open challenge, here we introduce the Condition Steiner Network (CSN) problem. In this setting, we are given a weighted undirected graph *G*, a set of *C* conditions and a set of *k* ≥ *C* demands, at least one per condition (note that we also cover the case of directed graphs, with similar results). The conditions are specified over a sequence of graphs *G_c_* defined over each condition, where vertices remain the same, but edges are allowed to change across conditions (notably, our results also hold when *G_c_* is defined with changing vertices rather than edges). Furthermore, demands are in the form of “connect node *u* to node *v* through a path of nodes that are present in condition *c*”. The goal is to find a minimum weight subgraph of *G* that satisfied all the demands.

We first show that it is NP-hard to find a solution that is better than the trivial one (obtained by solving the problem independently for each condition). This result extends to several types of connectivity problems and provides a theoretical lower bounds to the best-possible approximation guarantee that can be achieved in a multiple condition setting (Table 1). For instance, we can conclude that concomitantly solving the shortest path problem for a set of conditions is hard to approximate, and that the trivial solution (i.e., solving the problem to optimality in each condition) is, theoretically, the best that one can do. Another example, commonly used in PPI analysis, is the Prize-Collecting Steiner tree problem [7, 8]. Here, our results indicate that given a fixed input for this problem (i.e., a penalty function *p*(*v*) for each vertice), it is NP hard to solve it concomitantly in *C* conditions, such that the weight of the obtained solution is less than *C* times that of the optimal solution. Interestingly, a theoretical guarantee of
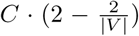
^1^ can be obtained by solving the problem independently for each time point

**Table 1.** Approximation bounds for the various Steiner Network Problems in their classic setting and condition setting.For the classic problems, we have indicated the papers in which the bounds are shown. For the condition problems, all the lower bounds are developed in the present work; all the upper bounds are the naive bounds obtained from the “union of shortest paths” heuristic, or from applying the best known approximation algorithm for the appropriate classic Steiner problem to each condition, then taking the union of those solutions.

| Problems | Classic |  | Condition |  |
| --- | --- | --- | --- | --- |
|  | Lower Bound | Upper Bound | Lower Bound(s) | Upper Bound(s) |
| Steiner Forest | 1.01 [16] | 2 [15] | $C - \epsilon, k - \epsilon$ | $2C, k$ |
| Directed Steiner Network | $k^{1/4-o(1)}$ [24] | $k^{1/2+\epsilon}$ [18, 19] | $C - \epsilon, k - \epsilon$ | $C \cdot k^{1/2+\epsilon}, k$ |
| Undirected Shortest Path | N/A | 1 | $C - \epsilon$ | $C$ |
| Directed Shortest Path | N/A | 1 | $C - \epsilon$ | $C$ |
| Steiner Tree | 1.01 [16] | 1.39 [14] | $C - \epsilon$ | $1.39C$ |
| Prize-Collecting Steiner Tree | 1.01 [16] | 1.97 [23] | $C - \epsilon$ | $1.97C$ |

**Table 2.** ILP solve times for some random instances generated by our random model.

| $\beta$ | $\gamma$ | $p$ | Time to solve |
| --- | --- | --- | --- |
| 10 | 1 | .25 | 1m ± 10s |
| 100 | 1 | .25 | 1m ± 10s |
| 10 | 1 | .75 | 1m ± 10s |
| 100 | 1 | .75 | 1m ± 10s |
| 10 | 10 | .25 | 9m ± 30s |
| 10 | 10 | .75 | 11m ± 30s |
| 100 | 10 | .25 | 12m ± 30s |
| 100 | 10 | .75 | 17m ± 2m |
| 10 | 100 | .75 | 2h 30m ± 12m |
| 100 | 100 | .75 | 4h ± 40m |

While these results provide a somewhat pessimistic view, they rely on the assumption that the network frames *G_c_* are arbitrary. In the last part of this paper, we show that for the specific case where the conditions can be ordered such that each condition is a subset of the next (namely, *G_c_* ⊆ *G_c_*′ for *c* ≤ *c*′) then the CSN problem can be reduced and solved as a standard connectivity problem with a single condition, leading to substantially better theoretical guarantees. Finally, we develop an Integer Linear Programming algorithm for the general CSN problem, and show that provided with real-world input (namely, the human PPI) it is capable of reaching an optimal solution in a reasonable amount of time.

## Introduction to Steiner Problems

The Steiner Tree problem, along with its many variants and generalizations, form a core family of NP-hard combinatorial optimization problems. Traditionally, the input to one of these problems is a single (usually weighted) graph, along with requirements about which nodes need to be connected in some way; the goal is to pick a minimum-weight subgraph satisfying the connectivity demands.

In this paper, we offer a *multi-condition* perspective; in our setting, multiple graphs over the same vertex set (which one can think of as an initial graph changing over a set of discrete conditions), are all given as input, and the goal is to pick a subgraph satisfying condition-sensitive connectivity requirements. Our study of this problem draws motivation and techniques from several lines of research, which we briefly summarize.

### Classic Steiner problems

A basic problem in graph theory is finding the *shortest path* between two nodes; this problem is efficiently solved using, for example, Dijkstra’s algorithm.

A natural extension of this is the Steiner Tree problem: given a weighted undirected graph *G* = (*V*,*E*) and a set of terminals *T* ⊆ *V*, find a minimum-weight subtree that connects all the nodes in *T* .A further generalization is Steiner Forest: given *G* = (*V*,*E*) and a set of demand pairs *D* ⊆ *V* × *V*, find a subgraph that connects each pair in *D*. Currently the best known approximation algorithms give a ratio of 1.39 for Steiner Tree [14] and 2 for Steiner Forest [15]. These problems are known to be NP-hard to approximate to within some small constant [16].

For directed graphs, we have the Directed Steiner Network (DSN) problem, in which we are given a weighted directed graph *G* = (*V*, *E*) and *k* demands (*a*_1_, *b*_1_),(*a_k_*, *b_k_*) ∈ *V* × *V*, and must find a minimum-weight sub-graph in which each *a_i_* has a path to *b_i_*. When *k* is fixed, DSN admits a polynomial-time exact algorithm [17]. For general *k*, the best known approximation algorithms have ratio *O*(*k*^1/2+*ϵ*^) for any fixed *ϵ* > 0 [18, 19]. On the complexity side, Dodis and Khanna [20] ruled out a polynomial-time *O*(2^log^1–*ϵ*^*n*^)-approximation for unless NP has quasipolynomial-time algorithms^[2]^. An important special case of DSN is Directed Steiner Tree, in which all demands have the form (*r*,*bi*) for some root node *r*. This problem has a *O*(*k^*ϵ*^*)-approximation scheme [21] and a lower bound of Ω(log^2–*ϵ*^ _*n*_) [22].

Finally, a Steiner variant that has found extensive use in computational biology is the Prize-Collecting Steiner Tree problem, in which the input contains a weighted undirected graph *G* = (*V*,*E*) and penalty function *p* : *V* → ℝ_≥0_; the goal is to find a subtree which simultaneously minimizes the weights of the edges in the tree and the penalties paid for nodes not included within the tree, i.e. cost(*T*) := Σ_*e*∈*T*_ *w*(*e*) + Σ_*v*∉*T*_ *p*(*v*). For this problem, an approximation algorithm with ratio 1.967 is known [23].

### Condition Steiner problems

In this paper, we generalize the Shortest Path, Steiner Tree, Steiner Forest, Directed Steiner Network, and Prize-Collecting Steiner Tree problems to the multi-condition setting. In this setting, we have a set of *conditions* [*C*] := {1,…,*C*}, and are given a graph for each condition.

Our main object of study is the natural generalization of Steiner Forest (in the undirected case) and Directed Steiner Network (in the directed case), which we call Condition Steiner Network:

#### Definition 1

(Condition Steiner Network (CSN)) *We are given the following inputs*:

1. *A sequence of undirected graphs G*_1_ = (*V*, *E*_1_), *G*_2_ = (*V*, *E_2_*),…, *G_C_* = (*V*, *_EC_*), *one for each* condition *c* ∈ [*C*]. *Each edge e in the underlying edge set E* := ⋃_*c*_ *E_c_ has a weight w*(*e*) ≥ 0.
2. *A set of k* connectivity demands 𝒟 ⊆ *V* × *V* × [*C*]. *We assume that for every c ∈ C there exists at least one demand and therefore that k* ≥ |*C*|.

*We call G* = (*V*, *E*) *the* underlying graph. *We say a subgraph H* ⊆ *G* satisfies *demand* (*a*,*b*,*c*) ∈ 𝒟 *iff H contains an a-b path P along which all edges exist in G_c_*. *The goal is to output a minimum-weight subgraph H* ⊆ *G that satisfies every demand in* 𝒟.

#### Definition 2

(Directed Condition Steiner Network (DCSN)) *This is the same as CSN except that all the edges are directed, and a demand* (*a*, *b*, *c*) *must be satisfied by a directed path from a to b in G_c_*.

We can also define the analogous generalizations of Shortest Path, (undirected) Steiner Tree, and Prize-Collecting Steiner Tree. We give hardness results and algorithms for these problems by demonstrating reductions to and from CSN and DCSN.

#### Definition 3

(Condition Shortest Path (CSP), Directed Condition Shortest Path (DCSP)) *These are the special cases of CSN and DCSN in which the demands are precisely* (*a*,*b*, 1),…, (*a*, *b*, *C*) *where a*,*b* ∈ *V are common source and target nodes*.

#### Definition 4

(Condition Steiner Tree (CST)) *We are given a sequence of undirected graphs G*_1_ = (*V*, *E*_1_),…,*G_C_* = (*V*, *E_C_*), *weights w*(⋅) ≥ 0 *on the edges*, *and sets of* terminal nodes *X*_1_,…, *X_C_* ⊆ *V*. *We say a subgraph H* ⊆ (*V*, ⋃_c_ *E_c_*) satisfies *the terminal set X_c_ iff the nodes in X_c_ are mutually reachable using edges in H that exist at condition c*. *The goal is to find a minimum-weight subgraph H that satisfies X_c_ for every c* ∈ [*C*].

#### Definition 5

(Condition Prize-Collecting Steiner Tree (CPCST)) *We are given a sequence of undirected graph G*_1_ = (*V*, *E*_1_), …, *G_C_* = (*V*, *E_C_*), *a weight w* (*e*) ≥ 0 *on each e* ∈ *E*, *and a penalty p*(*v*, *c*) ≥ 0 *for each v* ∈ *V*,*c* ∈ [*C*]. *The goal is to find a subtree T that minimizes* Σ_*e*∈*T*_ *w*(*e*) + Σ_*v*∈*T*,*c*∉[*C*]_ *p*(*v*, *c*).

Finally, in molecular biology applications, it is often the case that all the demands originate from a common root node. To capture this, we define the following special case of DCSN:

#### Definition 6

(Single-Source DCSN) *This is the special case of DCSN in which the demands are precisely* (*a*, *b*_1_, *c*_1_), (*a*, *b*_2_,*c*_2_),…, (*a*, *b_k_*, *c_k_*), *for some* root *a* ∈ *V*. *We can assume that c*_1_ ≤ *c*_2_ ≤ … ≤ *c_k_*.

It is also natural to consider variants of these problems in which nodes (rather than edges) vary across the conditions, or in which both nodes and edges vary. In Problem variants, we show that all three variants are in fact equivalent; thus we focus on the edge-based formulations.

## Our Results

In this work, we perform a systematic study of the condition Steiner problems defined above, from the standpoint of approximation algorithms—that is, algorithms that return subgraphs whose total weights are not too much greater than that of the optimal subgraph—as well as integer linear programming (ILP). Since all of the condition Steiner problems listed in the previous section turn out to be NP-hard (and in fact all of them except Shortest Path are hard even in the classic single-condition setting) we cannot hope for algorithms that are optimal and run in polynomial time.

First, in Hardness of condition Steiner problems, we show a series of strong negative results, starting with (directed and undirected) Condition Steiner Network:

### Theorem 1

(Main Theorem) *CSN and DCSN are NP-hard to approximate to a factor of C* – *ϵ as well as k* – *ϵ for every fixed k* ≥ 2 *and every constant *ϵ** > 0. *For DCSN*, *this holds even when the underlying graph is acyclic*.

Thus the best approximation ratio one can hope for is *C* or *k*; the latter upper bound is easily achieved by the trivial “union of shortest paths” algorithm: for each demand (*a*, *b*, *c*), compute the shortest *a-b* path at condition *c*; then take the union of these *k* paths. This contrasts with the classic Steiner network problems, which have nontrivial approximation algorithms and efficient fixed-parameter algorithms.

Next, we show similar hardness results for the other three condition Steiner problems. This is achieved by a series of simple reductions from CSN and DCSN.

### Theorem 2

*Condition Shortest Path*, *Directed Condition Shortest Path*, *Condition Steiner Tree*, *and Condition Prize-Collecting Steiner Tree are all NP-hard to approximate to a factor of C* – *ϵ for every fixed C* ≥ 2 *and ϵ* > 0.

Note that each of these condition Steiner problems can be naively approximated by applying the best known algorithm for the classic version of that problem in each graph in the input, then taking the union of all those subgraphs. If the corresponding classic Steiner problem can be approximated to a factor of *α*, then this process gives an *α* ⋅ *C*-approximation for the condition version. Thus using known constant-factor approximation algorithms, each of the condition problems in Theorem 2 has an *O*(*C*)-approximation algorithm. Our result shows that in the worst case, one cannot do much better.

While these results provide a somewhat pessimistic view, the proofs rely on the assumption that the edge sets in the input networks (that is, *E*_1_,…, *E_C_*) do not necessarily bear any relationship to one another. In Monotonic special cases, we move beyond this worst-case assumption by studying a broad class of special cases in which the conditions are *monotonic*: if an edge *e* exists in some graph *G_c_*, then it exists in all the subsequent graphs *G_c′_*, *c*′ ≥ *c*. In other words, each graph in the input is a subgraph of the next. For these problems, we prove the following two theorems:

### Theorem 3

*Monotonic CSN has a polynomial-time O*(log *k*)-*approximation algorithm*. *It has no* Ω(log log*n*)-*approximation algorithm unless NP* ∈ DTIME(*n*^log loglog *n*^).

In the directed case, for monotonic DCSN with a single source (that is, every demand is of the form (*r*, *b*, *c*) for a common root node *r*), we show the following:

### Theorem 4

*Monotonic Single-Source DCSN has a polynomial-time O*(*k*^*ϵ*^)-*approximation algorithm for every ϵ* > 0. *It has no* Ω(log^2–*ϵ*^ *n*)-*approximation algorithm unless* NP ∈ ZPTIME(*n*^polylog(*n*)^).

These bounds are proved via approximation-preserving reductions to and from classic Steiner problems, namely Priority Steiner Tree and Directed Steiner Tree. Conceptually, this demonstrates that imposing the monotonicity requirement makes the condition Steiner problems much closer to their classic counterparts, allowing us to obtain algorithms with substantially better approximation guarantees.

Finally in Application to protein-protein interaction networks, we show how to model various condition Steiner problems as integer linear programs (ILPs). In experiments on real-world inputs derived from the human PPI network, we find that these ILPs are capable of reaching optimal solutions in a reasonable amount of time.

Table 1 summarizes our results, emphasizing how the known upper and lower bounds change when going from the classic Steiner setting to the condition Steiner setting.

## Preliminaries

Note that the formulations of CSN and DCSN in the Introduction involved a fixed vertex set; only the edges change over the conditions. It is also natural to formulate the condition Steiner network problem with nodes changing over condition, or both nodes and edges. However by the following proposition, it is no loss of generality to discuss only the edge-condition variant.

### Proposition 1

*The edge*, *node*, *and node-and-edge variants of CSN are mutually polynomial-time reducible via strict reductions (i.e*. *preserving the approximation ratio exactly). Similarly all three variants of DCSN are mutually strictly reducible*.

We defer the precise definitions of the other two variants, as well as the proof of this proposition, to Problem variants.

Next we state the Label Cover problem, which is the starting point of one of our reductions to CSN.

### Definition 7

(Label Cover (LC)) *An instance of this problem consists of a bipartite graph G* = (*U*, *V*, *E*) *and a set of possible labels* Σ. *The input also includes, for each edge* (*u*,*v*) ∈ *E*, projection functions
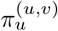
: Σ → *C and*
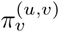
: Σ → *C*, *where C is a common set of colors*;
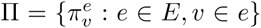
*is the set of all such functions*. *A labeling of G is a function ϕ* : *U* ∪ *V* → Σ *assigning each node a label*. *We say a labeling ϕ* satisfies *an edge* (*u*,*v*) ∈ *E*, *or* (*u*, *v*) *is consistent under ϕ*, *iff*
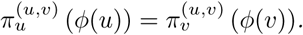
*The task is to find a labeling that satisfies as many edges as possible*.

This slightly generalizes the original definition in [25]. It has the following gap hardness, which follows by combining the PCP theorem [26] with Raz’s parallel repetition theorem [27].

### Theorem 5

*For every ϵ* > 0, *there is a constant* |Σ| *such that the following promise problem is NP-hard: Given a Label Cover instance* (*G*, Σ, Π), *distinguish between the following cases*:

- *(YES instance) There exists a* total labeling *of G; i.e*. *a labeling that satisfies every edge*.
- *(NO instance) There does not exist a labeling of G that satisfies more than *ϵ**|*E*| *edges*.

In Hardness of condition Steiner problems, we use Label Cover to show (2 – *ϵ*)-hardness for 2-CSN and 2-DCSN; that is, when there are only two demands. To prove our main result however, we will actually need a generalization of Label Cover to partite hypergraphs, called *k*-Partite Hypergraph Label Cover. Out of space considerations we defer the statement of this problem and its gap hardness to Inapproximability for general *C* and *k*, where the (2 –*ϵ*)-hardness result is generalized to show (*C* – *ϵ*)-hardness and (*k* – *ϵ*)-hardness for general number of conditions *C* and demands *k*.

## Hardness of condition Steiner problems

### Overview of the reduction

Here we outline our strategy for reducing Label Cover to the condition Steiner problems. First, we reduce to the CSN problem restricted to having only *C* = 2 conditions and *k* = 2 demands; we call this problem 2-CSN. The directed problem 2-DCSN is defined analogously. Later, we obtain similar hardness for CSN with more conditions or demands by using the same ideas, but reducing from *k*-Partite Hypergraph Label Cover.

Consider the nodes *u*_1_,…, *u*_|*U*|_ on the “left” side of the LC instance. We build, for each *u_i_*, a gadget (which is a small sub-graph in the Steiner instance) consisting of multiple parallel directed paths from a source to a sink—one path for each possible label for *u_i_*. We then chain together these gadgets, so that the sink of *u*_1_’s gadget is the source of *u*_2_’s gadget, and so forth. Finally we create a connectivity demand from the source of *u*_1_’s gadget to the sink of *u*|*U*|’s gadget, so that a solution to the Steiner instance must have a path from *u*_1_’s gadget, through all the other gadgets, and finally ending at *u*_|*u*|_’s gadget. This path, depending on which of the parallel paths it takes through each gadget, induces a labeling of the left side of the Label Cover instance. We build an analogous chain of gadgets for the nodes on the right side of the Label Cover instance.

The last piece of the construction is to ensure that the Steiner instance has a low-cost solution if and only if the Label Cover instance has a consistent labeling. This is accomplished by setting all the *u_i_* gadgets to exist only at condition 1 (i.e. in frame *G*_1_), setting the *V_j_* gadgets to exist only in *G*_2_, and then merging certain edges from the *u_i_*-gadgets with edges from the *v_j_*-gadgets, replacing them with a single, shared edge that exists in both frames. Intuitively, the edges we merge are from paths that correspond to labels that satisfy the Label Cover edge constraints. The result is that a YES instance of Label Cover (i.e. one with a total labeling) will enable a high degree of overlap between paths in the Steiner instance, so that there is a very low-cost solution. On the other hand, a NO instance of LC will not result in much overlap between the Steiner gadgets, so every solution will be costly.

Let us define some of the building blocks of the reduction we just sketched:

- A *bundle* is a graph gadget consisting of a source node *b*_1_, sink node *b*_2_, and parallel, disjoint *strands* (defined shortly) from *b*_1_ to *b*_2_.
- A *chain* of bundles is a sequence of bundles, with the sink of one bundle serving as the source of another.
- A *simple strand* is a directed path of the form *b*_1_ → *c*_1_ → *C*_2_ → *b*_2_.
- In a simple strand, we say that (*c*_1_, *c*_2_) is the *contact edge*. Contact edges have weight 1; all other edges in our construction have zero weight.
- More generally, a *strand* can be made more complicated, by replacing a contact edge with another bundle (or even a chain of them). In this way, bundles can be nested, as shown in Figure 1. **Figure 1.**
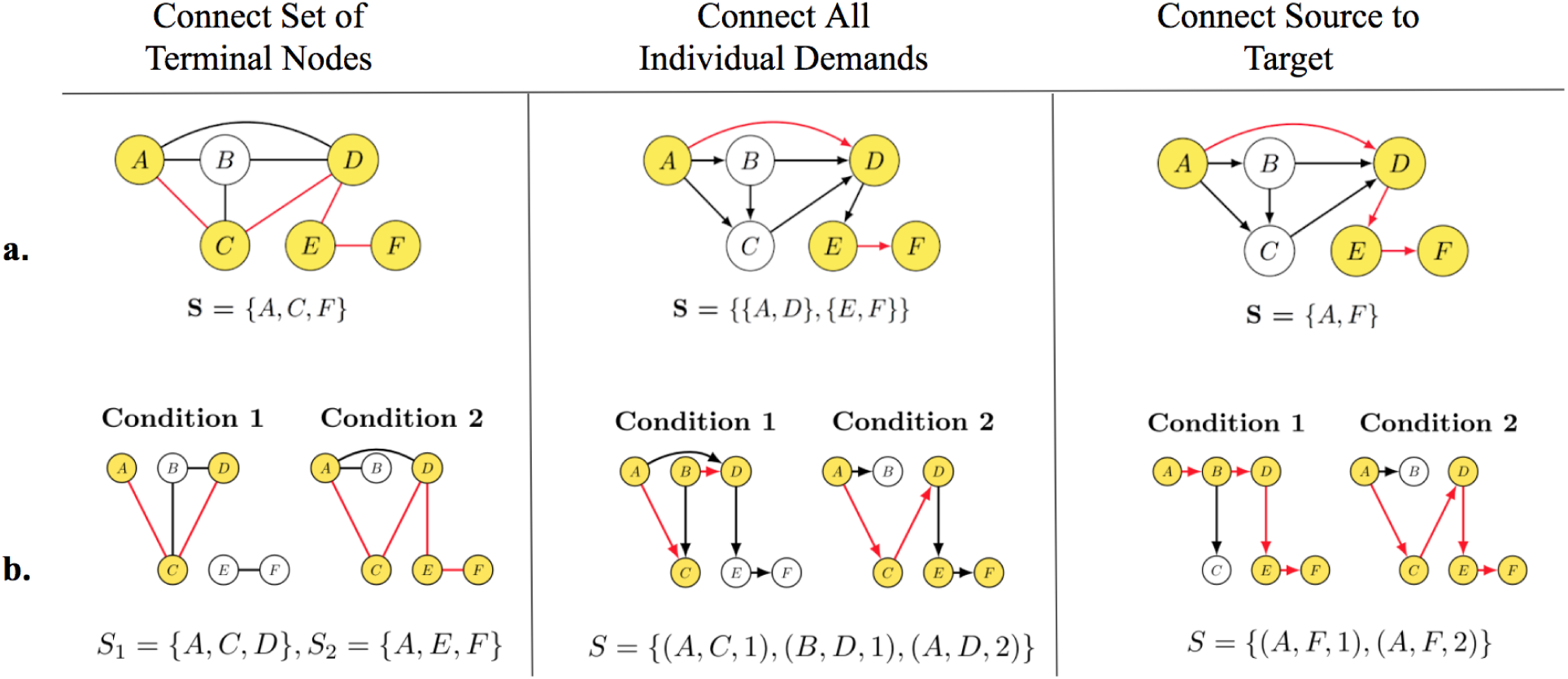
Examples of well studied network problems (a), and their corresponding extension with multiple conditions (b). The problems shown are: Undirected Steiner Tree, Directed Steiner Network, and Shortest Path, respectively. Yellow nodes and red edges correspond to nodes and edges used in the optimal solutions for the corresponding instances. **Figure 2.**
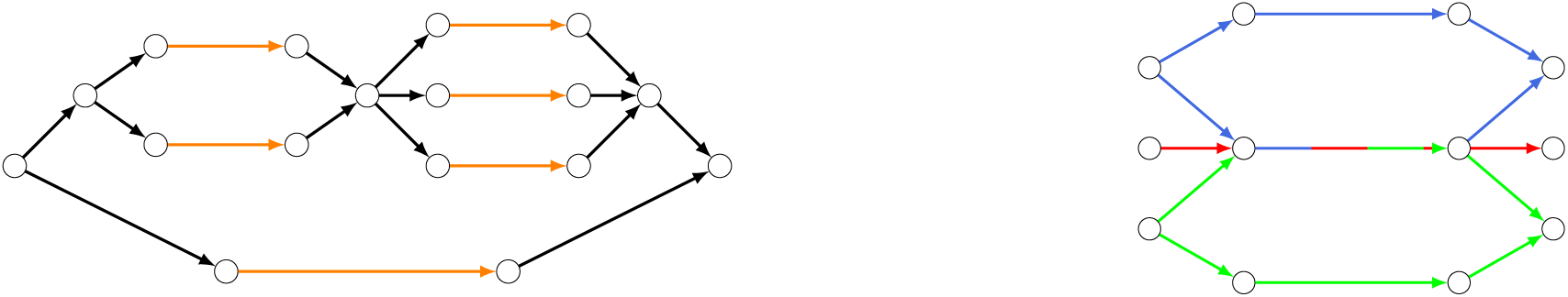
(Left) A bundle whose upper strand is a chain of two bundles; the lower strand is a simple strand. Contact edges are orange. (Right) Three bundles (blue, green, red indicate different conditions), with one strand from each merged together. **Figure 3.**
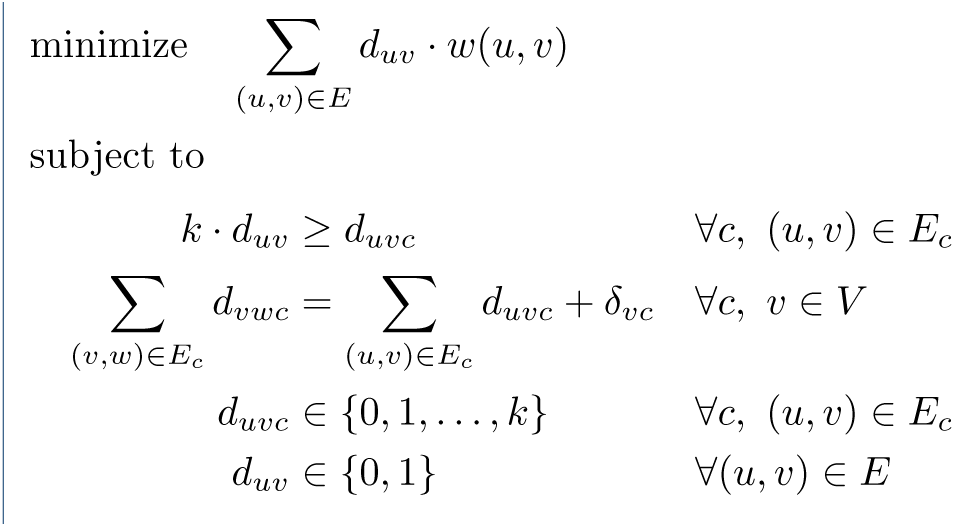
Integer Linear Program for Single-Source Conditon Steiner Network. *S_vc_* = number of demands at condition c in which *v* is a source, minus the number in which *v* is a target.
- We can *merge* two or more simple strands from different bundles by setting their contact edges to be the same edge, and making that edge existent at the union of all conditions when the original edges existed (Figure 1).

Before formally giving the reduction, we illustrate a simple example of its construction.

#### Example 1

*Consider a toy Label Cover instance whose bipartite graph is a single edge, label set is* Σ = {1, 2}, *and projection functions are shown*:

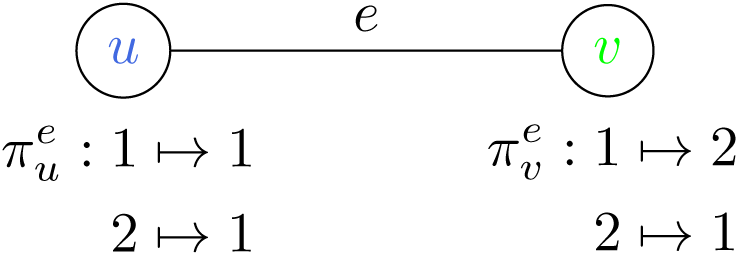

*Our reduction outputs this corresponding 2-CSN instance:*

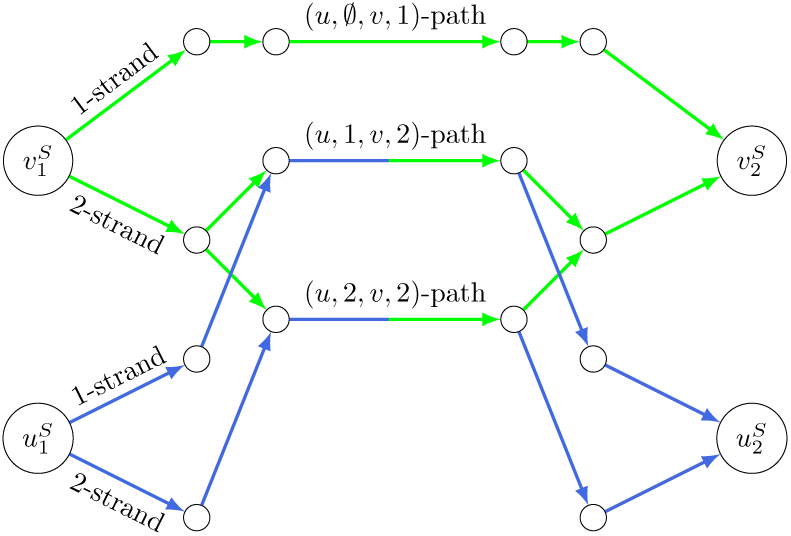

*G*_1_ *comprises the set of blue edges; G*_2_ *is green*. *The demands are*
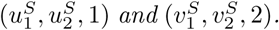
*The gadget for the Label Cover node u (the blue sub-graph) consists of two strands, one for each possible label. In the v-gadget (green sub-graph)*, *the strand corresponding to a labeling of ‘2’ branches further*, *with one simple strand for each agreeing labeling of u*. *Finally*, *strands (more precisely*, *their contact edges) whose labels map to the same color are merged*.

*The input is a YES instance of Label Cover whose optimal labelings (u gets either label 1 or 2*, *v gets label 2) correspond to 2-CSN solutions of cost 1 (both gadgets traverse the* (*u*, *1*,*v*, *2*)-*path*, *or both traverse the* (*u*, *2*, *v*, *2*)-*path*). *If this were a NO instance and edge e could not be satisfied*, *then the resulting 2-CSN gadgets would have no overlap*.

### Inapproximability for two demands

We now formalize the reduction in the case of two conditions and two demands; later, we extend this to general *C* and *k*.

#### Theorem 6

*2-CSN and 2-DCSN are NP-hard to approximate to within a factor of* 2 – *ϵ for every constant ϵ* > 0. *For 2-DCSN*, *this holds even when the underlying graph is acyclic*.

#### Proof

We describe a reduction from Label Cover (LC) to 2-DCSN with an acyclic graph. Given the LC instance (*G* = (*U*,*V*,*E*), Σ, Π), construct a 2-DCSN instance (𝒢 = (*G*_1_,*G*_2_), along with two connectivity demands) as follows. Create nodes
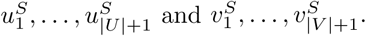
Let there be a bundle from each
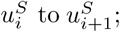
we call this the *u_i_-bundle*, since a choice of path from
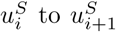
in 𝒢 will indicate a labeling of *u_i_* in *G*.

The *u_i_*-bundle has a strand for each possible label *ℓ* ∈ Σ. Each of these *ℓ-strands* consists of a chain of bundles—one for each edge (*u_i_*, *v*) ∈ *E*. Finally, each such (*u_i_*, *v*)-*bundle* has a simple strand for each label *r* ∈ Σ such that
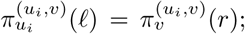
call this the (*u_i_*, *ℓ*, *v*, *r*)-*path*. In other words, there is ultimately a simple strand for each possible labeling of *u_i_*’s neighbor *v* such that the two nodes are in agreement under their mutual edge constraint. If there are no such consistent labels *r*, then the (*u_i_*, *v*)-bundle consists of just one simple strand, which is not associated with any *r*. Note that every minimal
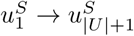
path (that is, one that proceeds from one bundle to the next) has total weight exactly |*E*|.

Similarly, create a *V_i_*-bundle from each
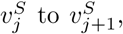
whose *r*-strands (for *r* ∈ Σ) are each a chain of bundles, one for each (*u*, *v_j_*) ∈ *E*. Each (*u*, *v_j_*)-bundle has a (*u*, *ℓ*, *v_j_*, *r*)-path for each agreeing labeling *ℓ* of the neighbor *u*, or a simple strand if there are no such labelings.

Set all the edges in the *u_i_*-bundles to exist in *G*_1_ only. Similarly the *v_j_*-bundles exist solely in *G*_2_. Now, for each (*u*, *ℓ*, *v*, *r*)-path in *G*_1_, merge it with the (*u*, *ℓ*, *v*, *r*)-path in *G*_2_, if it exists. The demands are
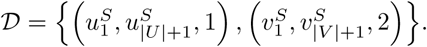

We now analyze the reduction. The main idea is that any
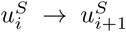
path induces a labeling of *u_i_*; thus the demand
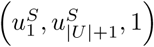
ensures that any 2-DCSN solution indicates a labeling of all of *U*. Similarly,
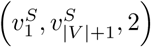
forces an induced labeling of *V*. In the case of a YES instance of Label Cover, these two connectivity demands can be satisfied by taking two paths with a large amount of overlap, resulting in a low-cost 2-DCSN solution. In contrast when we start with a NO instance of Label Cover, any two paths we can choose to satisfy the 2-DCSN demands will be almost completely disjoint, resulting in a costly solution. We now fill in the details.

Suppose the Label Cover instance is a YES instance, so that there exists a labeling
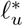
to each *u* ∈ *U*, and
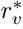
to each *v* ∈ *V*, such that for all edges (*u*, *v*) ∈ *E*,
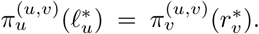
The following is an optimal solution ℋ^∗^ to the constructed 2-DCSN instance:

- To satisfy the demand at condition 1, for each *u*-bundle, take a path through the
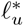
-strand. In particular for each (*u*, *v*)-bundle in that strand, traverse the
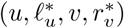
path.
- To satisfy the demand at condition 2, for each *v*-bundle, take a path through the
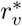
-strand. In particular for each (*u*,*v*)-bundle in that strand, traverse the
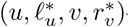
-path.

In tallying the total edge cost, ℋ^∗^ ∩ *G*_1_ (i.e. the subgraph at condition 1) incurs a cost of |*E*|, since one contact edge in 𝒢 is encountered for each edge in *G*. ℋ^∗^ ∩ *G*_2_ accounts for no additional cost, since all contact edges correspond to a label which agrees with some neighbor’s label, and hence were merged with the agreeing contact edge in ℋ^∗^ ∩ *G*_1_. Clearly a solution of cost |*E* | is the best possible, since every
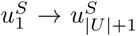
path in *G*_1_ (and every
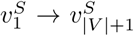
path in *G*_2_) contains at least |*E*| contact edges.

Conversely suppose we started with a NO instance of Label Cover, so that for any labeling
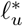
to *u* and
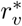
to *v*, for at least (1 – *ϵ*)|*E*| of the edges (*u*,*v*) ∈ *E*, we have
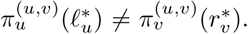
By definition, any solution to the constructed 2-DCSN instance contains a simple
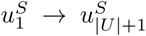
path *P*_1_ ∈ *G*_1_ and a simple
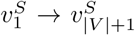
path *P*_2_ ∈ *G*_2_. *P*_1_ alone incurs a cost of exactly |*E*|, since one contact edge in 𝒢 is traversed for each edge in *G*. However, *P*_1_ and *P*_2_ share at most *ϵ*|*E*| contact edges (otherwise, by the merging process, this implies that more than *ϵ*|*E*| edges could be consistently labeled, which is a contradiction). Thus the solution has a total cost of at least (2 – *ϵ*)|*E*|.

The underlying directed graph we constructed is acyclic, as every edge points “to the right” as in Example 1. It follows from the gap between the YES and NO cases that 2-DCSN is NP-hard to approximate to within a factor of 2 – *ϵ* for every *ϵ* > 0, even on acyclic graphs. Finally, note that the same analysis holds for 2-CSN, by simply making every edge undirected; however in this case the graph is clearly not acyclic. □

### Inapproximability for general C and k

#### Theorem 1

(Main Theorem) *CSN and DCSN are NP-hard to approximate to a factor of C* – *ϵ as well as k* – *ϵ for every fixed k* ≥ 2 *and every constant ϵ* > 0. *For DCSN*, *this holds even when the underlying graph is acyclic*.

#### Proof

We perform a reduction from *k*-Partite Hypergraph Label Cover, a generalization of Label Cover to hypergraphs, to CSN, or DCSN with an acyclic graph. Using the same ideas as in the *C* = *k* = 2 case, we design *k* demands composed of parallel paths corresponding to labelings, and merge edges so that a good global labeling corresponds to lots of overlaps between those paths. The full proof is left to Inapproximability for general *C* and *k*.□

Note that a *k*-approximation algorithm is to simply choose ℋ = ⋃_*c*_ *p̃_c_*, where *p̃_c_* is the shortest *a_c_* → *b_c_* path in *G_c_*. Thus by Theorem 1, essentially no better approximation is possible in terms of *k* alone. In contrast, most classic Steiner problems have good approximation algorithms, or are even exactly solvable for constant *k*.

### Inapproximability for Steiner variants

We take advantage of our previous hardness of approximation results in Theorem 1 and show, via a series of reductions, that CSP, CSN, and CPCST are also hard to approximate.

#### Theorem 2

*Condition Shortest Path*, *Directed Condition Shortest Path*, *Condition Steiner Tree*, *and Condition Prize-Collecting Steiner Tree are all NP-hard to approximate to a factor of C* – *ϵ for every fixed C* ≥ 2 *and ϵ* > 0.

#### Proof

We first reduce from CSN to CSP (and DCSN to DCSP). Suppose we are given an instance of CSN with graph sequence 𝒢 = (*G*_1_,…, *G_c_*), underlying graph *G* = (*V*,*E*), and demands 𝓓 = {(*a_i_*,*b_i_*,*c_i_*) : *i* ∈ [*k*]}. We build a new instance
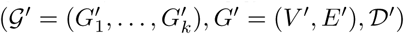
as follows.

Initialize *G*′ to *G*. Add to *G*′ the new nodes *a* and *b*, which exist at all conditions/in all frames
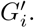
For all *e* ∈ *E* and *i* ∈ [*k*], if *e* ∈ *G_Ci_*, then let e exist in
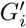
as well. For each (*a_i_*, *b_i_*, *c_i_*) ( 𝓓,

1. Create new nodes *x_i_*, *y_i_*. Create zero-weight edges (*a*, *x_i_*), (*x_i_*,*a_i_*), (*b_i_*, *_i_*), and (*y_i_*,*b*).
2. Let (*a*,*x_i_*), (*x_i_*,*a_i_*), (*b_i_*,*y_i_*), and (*y_i_*,*b*) exist only in frame
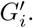

Lastly, the demands are 𝓓′ = {(*a*, *b*, *i*) : *i* ∈ [*k*]}.

Given a solution *H*′ ⊆ *G*′ containing an *a* → *b* path at every condition *i* ∈ [*k*], we can simply exclude nodes *a*, *b*, {*x_i_*}, and {*y_i_*} to obtain a solution *H* ⊆ *G* to the original instance, which contains an *a_i_* → *b_i_* path in *G_Ci_* for all *i* ∈ [*k*], and has the same cost. The converse is also true by including these nodes.

Observe that essentially the same procedure shows that DCSN reduces to DCSP; simply ensure that the edges added by the reduction are directed rather than undirected.

Next, we reduce CSP to CST. Suppose we are given an instance of CSP with graph sequence 𝓖 = (*G*_1_,…, *G_c_*), underlying graph *G* = (*V*, *E*), and demands 𝓓 = {(*a*, *b*, *i*) : *i* ∈ [*C*]}. We build a new instance of CST as follows:

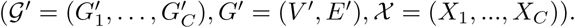

Set *𝓖*′ to 𝓖, and *G*^′^ to *G*. Take the set of terminals in each condition to be *X_i_* = {*a*, *b*}. We note that a solution *H*^′^ ⊆ *G*^′^ to the CST instance is trivially a solution the CSP instance with the same cost, and vice-versa.

Finally, we reduce CST to CPCST. We do this by making an appropriate assignment of the penalties *p*(*v*,*c*). Suppose we are given an instance of CST with graph sequence 𝓖 = (*G*_1_,…,*G_C_*), underlying graph *G* = (*V*, *E*), and terminal sets *χ* = (*X*_1_,…,*X_C_*). We build a new instance of CPCST,
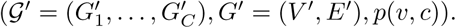
In particular, set 𝓖′ to 𝓖, and *G*′ to *G*. Set *p*(*v*, *c*) as follows:

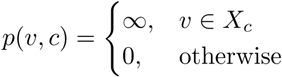

Consider any solution *H* ⊆ *G* to the original CST instance. Since *H* spans the terminals *X*_1_,…, *X_c_* (thus avoiding any infinite penalties), and since the nonterminal vertices have zero cost, the overall cost of *H* remains the same cost in the constructed CPCST instance. Conversely, suppose we are given a solution *H*′ ⊆ *G*′ to the constructed CPCST instance. If the cost of *H*′ is ∞, then there exists no solution that can span all the *X_c_*’s simultaneously, and thus no solution exists for the CST instance. On the other hand if H^’^ has finite cost, then *H*^′^ is also a solution for the CST instance, with the same cost.

To summarize: in the first reduction from CSN to CSP, the number of demands, *k*, in the CSN instance is the same as the number of the conditions, *C*, in the CSP instance; we conclude that CSP is NP-hard to approximate to a factor of *C* – *ϵ* for every fixed *C* ≥ 2 and *ϵ* > 0. Since *C* remains the same in the two subsequent reductions, we also have that CST and CPCST are NP-hard to approximate to a factor of *C* – *ϵ*. □

## Monotonic special cases

In light of the strong lower bounds in the previous theorems, in this section we consider more tractable special cases of the condition Steiner problems. A natural restriction is that the changes over conditions are *monotonic*:

### Definition 8

(Monotonic {CSN, DCSN, CSP, DCSP, CST, CPCST}) *In this special case (of any of the condition Steiner problems)*, *we have that for each e* ∈ *E and c* ∈ [*C*], *if e* ∈ *G_c_*, *then e* ∈ *G_c′_*≥ *for all c*′ ≥ *c*.

We now examine the effect of monotonicity on the complexity of the condition Steiner problems.

### Monotonicity in the undirected case

In the undirected case, we show that monotonicity has a simple effect: it makes CSN equivalent to the following well-studied problem:

#### Definition 9

(Priority Steiner Tree [28]) *The input is a weighted undirected multigraph G* = (*V*,*E*,*w*), *a* priority level *p*(*e*) *for each e* ∈ *E*, *and a set of k demands* (*a_i_*,*b_i_*), *each with priority p*(*a_i_*,*b_i_*). *The output is a minimum-weight forest F* ⊆ *G that contains, between each a_i_ and bi*, *a path in which every edge e has priority p*(*e*) ≤ *p*(*a_i_*, *b_i_*).

Priority Steiner Tree was introduced by Charikar, Naor, and Schieber [28], who gave a *O*(log *k*) approximation algorithm. Moreover, it cannot be approximated to within a factor of Ω(loglog *n*) assuming NP ∉ DTIME(*n*^log log log *n*^) [29]. We now show that the same bounds apply to Monotonic CSN.

#### Lemma 1

*Priority Steiner Tree and Monotonic CSN have the same approximability*.

#### Proof

We transform an instance of Priority Steiner Tree into an instance of Monotonic CSN as follows: the set of priorities becomes the set of conditions; if an edge *e* has priority *p*(*e*), it now exists at all conditions *t* ≥ *p*(*e*); if a demand (*a_i_*,*b_i_*) has priority *p*(*a_i_*,*b_i_*), it now becomes (*a_i_*, *b_i_*,*p*(*a_i_*, *b_i_*)). If there are parallel multiedges, break up each such edge into two edges of half the original weight, joined by a new node. Given a solution *H* ⊆ *G* to this CSN instance, contracting any edges that were originally multiedges gives a Priority Steiner Tree solution of the same cost. This reduction also works in the opposite direction (in this case there are no multiedges), which shows the equivalence. □

Furthermore, the *O*(log *k*) upper bound applies to CST (We note that Monotonic CSP admits a trivial algorithm, namely take the subgraph induced by running Djikstra’s Algorithm on *G*_1_).

#### Lemma 2

*If Monotonic CSN can be approximated to a factor of f* (*k*) *for some function f*, *then Monotonic CST can also be approximated to within f* (*k*).

#### Proof

We now show a reduction from CST to CSN. Suppose we are given a CST instance on graphs 𝓖 = (*G*_1_,…, *G_C_*) and terminal sets *χ* = (*X*_1_,…, *X_C_*). Our CSN instance has precisely the same graphs, and has the following demands: for each terminal set *X_c_*, pick any terminal *a* ∈ *X_c_* and create a demand (*a*, *b*, *c*) for each *b* ≠ *a* ∈ *X_c_*. A solution to the original CST instance is a solution to the constructed CSN instance with the same cost, and vice-versa; moreover, if the CST instance is monotonic, then so is the constructed CSN instance. Observe that if the total number of CST terminals is *k*, then the number of constructed demands is *k* – *C*, and therefore an *f* (*k*)-approximation for CSN implies an *f* (*k* – *C*) ≤ *f* (*k*)-approximation for CST, as required.□

### Monotonicity in the directed case

In the directed case, we give an approximation-preserving reduction from a single-source special case of DCSN to the Directed Steiner Tree (DST) problem, then apply a known algorithm for DST. Recall the definition of Single-Source DCSN:

#### Definition 6

(Single-Source DCSN) *This is the special case of DCSN in which the demands are precisely* (*a*, *b*_1_, *c*_1_), (*a*, *b*_2_,*c*_2_),…, (*a*, *b_k_*, *c_k_*), *for some* root *a* ∈ *V*. *We can assume that c*_1_ ≤ *c*_2_ ≤ ⋯ ≤ *c_k_*.

#### Lemma 3

*Monotonic Single-Source DCSN and Directed Steiner Tree have the same approximability*.

For the remainder of this section, we refer to Monotonic Single-Source DCSN as simply DCSN. Towards proving the theorem, we now describe a reduction from DCSN to DST. Given a DCSN instance (*G*_1_ = (*V*,*E*_1_),*G*_2_ = (*V*,*E*_2_),…,*G_C_* = (*V*,*E_C_*),*D*) with underlying graph 𝓖 = (*V*, *E*), we construct a DST instance (*G*′ = (*V*′, *E*′),*D*′) as follows:

- *G*′ contains a vertex *v^i^* for each *v* ∈ *V* and each *i* ∈ [*k*]. It contains an edge (*u^i^*, *v^i^*) with weight *w*(*u*, *v*) for each (*u*,*v*) ∈ *E_i_*. Additionally, it contains a zero-weight edge (*v^i^*,*v*^*i*+1^) for each *v* ∈ *V* and each *i* ∈ [*k*].
- *D*′ contains a demand
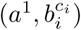
for each (*a*, *b_i_*, *c_i_*) ∈ 𝓓. Now consider the DST instance (*G*′,*D*′).

#### Lemma 4

*If the DCSN instance* (*G*_1_,…,*G_C_*, *D*) *has a solution of cost C*^∗^, *then the constructed DST instance* (*G*′,*D*′) *has a solution of cost at most C*^∗^.

#### Proof

Let 𝓗 ⊆ 𝓖 be a DCSN solution having cost *C*^∗^. For any edge (*u*,*v*) ∈ *E*(𝓗), define the *earliest necessary condition* of (*u*, *v*) to be the minimum *c_i_* such that removing (*u*, *v*) would cause 𝓗 not to satisfy demand (*a*, *b_i_*, *c_i_*).

#### Claim 1

*There exists a solution* 𝓒 ⊆ 𝓗 *that is a directed tree and has cost at most C*^∗^. *Moreover for every path P_i_ in* 𝓒 *from the root a to some target b_i_*, *as we traverse P_i_ from a to b_i_*, *the earliest necessary conditions of the edges are non-decreasing*.

*Proof of Claim 1* Consider a partition of 𝓗 into edge-disjoint sub-graphs 𝓗_1_,…, 𝓗_*k*_, where 𝓗_*i*_ is the subgraph whose edges have earliest necessary condition *c_i_*. Clearly each 𝓗_*i*_ is a single component.

If there is a directed cycle or parallel paths in the first sub-graph 𝓗_1_, then there is an edge *e* ∈ *E*(𝓗_1_) whose removal does not cause 𝓗_1_ to satisfy fewer demands at condition *c*_1_. Moreover by monotonicity, removing e also does not cause 𝓗 to satisfy fewer demands at any future conditions. Hence there exists a directed tree 𝓣_1_ ⊆ 𝓗_1_ such that 𝓣_1_ ∪ 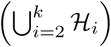 has cost at most *C*^∗^ and still satisfies 𝓣.

Now suppose by induction that for some *j* ∈ [*k* – 1], 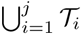 is a tree such that 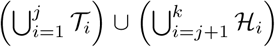 has cost at most *C*^∗^ and satisfies 𝓓. Consider the partial solution 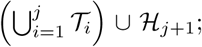 if this sub-graph is not a directed tree, then there must be an edge (*u*,*v*) ∈ *E*(𝓗_*j*+1_) such that *v* has another in-edge in the sub-graph. However by monotonicity, (*u*, *v*) does not help satisfy any new demands, as *v* is already reached by some other path from the root. Hence by removing all such redundant edges, we have 𝓣_*j*+_1__ ⊆ 𝓗_*j*+1_ such that 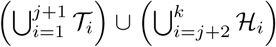 has cost at most *C*^∗^ and satisfies 𝓓, which completes the inductive step.

We conclude that 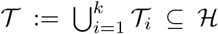 is a tree of cost at most *C*^∗^ satisfying 𝓓. Observe also that by construction, 𝓣 has the property that if we traverse any *a* → *b_i_* path, the earliest necessary conditions of the edges never decrease. □

Now let 𝓣 be the DCSN solution guaranteed to exist by *Claim 1*. Consider the sub-graph *H*′ ⊆ *G*′ formed by adding, for each (*u*, *v*) ∈ *E*(𝓣), the edge (*u^c^*,*v^c^*) ∈ *E*′ where *c* is the earliest necessary condition of (*u*, *v*) in *E*(𝓗). In addition, add all the free edges (*v^i^*, *v*^*i*+1^). Since *w*(*u^c^*,*v^c^*) = *w*(*u*,*v*) by construction, cost(*H*′) ≤ cost(𝓣) ≤ *C*^∗^.

To see that *H*′ is a valid solution, consider any demand 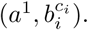 Recall that 𝓣 has a unique *a* → *b_i_* path *P_i_* along which the earliest necessary conditions are nondecreasing. We added to *H*′ each of these edges at the level corresponding to its earliest necessary condition; moreover, whenever there are adjacent edges (*u*, *v*), (*v*, *x*) ∈ *P_i_* with earliest necessary conditions *c* and *c*′ ≥ *c* respectively, there exist in *H*′ free edges (*v^t^*, *v*^*c*+1^),…, (*v*^*c*–1^,*v*^*c*′^). Thus *H*′ contains an 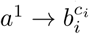 path, which completes the proof. □

#### Lemma 5

*If the constructed DST instance* (*G*′, *D*′) *has a solution of cost C*^∗^, *then the original DCSN instance* (*G*_1_,…, *G_C_*, 𝓓) *has a solution of cost at most C*^∗^.

#### Proof

First note that any DST solution ought to be a tree; let *T*′ ⊆ *G*′ be such a solution of cost *C*. For each (*u*, *v*) ∈ *G*, *T*′ might as well use at most one edge of the form (*u^i^*, *v^i^*), since if it uses more, it can be improved by using only the one with minimum *i*, then taking the free edges (*v^i^*,*v*^*i*+1^) as needed. We create a DCSN solution 𝓣 ⊆ 𝓖 as follows: for each (*u^i^*,*v^i^*) ∈ *E*(*T*′), add (*u*, *v*) to 𝓣. Since *w*(*u*,*v*) = *w*(*u^i^*,*v^i^*) by design, we have cost(𝓣) ≤ cost(*T*′) ≤ *C*. Finally, since each 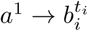 path in *G*′ has a corresponding path in 𝓖 by construction, 𝓣 satisfies all the demands. □

Lemma 3 follows from Lemma 4 and Lemma 5. Finally we can obtain the main result of this subsection:

#### Theorem 4

*Monotonic Single-Source DCSN has a polynomial-time O*(*k*^*ϵ*^)-*approximation algorithm for every ϵ* > 0. *It has no* Ω(log^2–*ϵ*^ *n*)-*approximation algorithm unless* NP ∈ ZPTIME(*n*^polylog(*n*)^).

#### Proof

The upper bound follows by composing the reduction (from Monotonic Single-Source DCSN to Directed Steiner Tree) with the algorithm of Charikar et al. [21] for Directed Steiner Tree, which achieves ratio *O*(*k*^*ϵ*^) for every *ϵ* > 0. More precisely they give a *i*^2^(*i* – 1)*k*^1/*i*^-approximation for any integer *i* ≥ 1, in time *O*(*n^i^ k*^2*i*^). The lower bound follows by composing the reduction (in the opposite direction) with a hardness result of Halperin and Krauthgamer [22], who show the same bound for Directed Steiner Tree. □

In Explicit algorithm for Monotonic Single-Source DCSN, we show how to modify the algorithm of Charikar et al. to arrive at a simple, explicit algorithm for Monotonic Single-Source DCSN achieving the same guarantee.

## Application to protein-protein interaction networks

Methods such as Directed Condition Steiner Network can be key in identifying underlying structure in biological processes. As a result, it is important to assess the runtime feasibility of solving for an optimal solution. We show via simulation on human protein-protein interaction networks, that our algorithm on single-source instances is able to quickly and accurately infer maximum likelihood subgraphs for a certain biological process.

### Building the protein-protein interaction network

We represent the human PPI network as a weighted directed graph, where proteins serve as nodes, and interactions serve as edges. The network was formed by aggregating information from four sources of interaction data, including Netpath [30], Phosphosite [31], HPRD [32], and InWeb [33], altogether, covering 16222 nodes and 437888 edges. Edge directions are assigned where these annotations were available (primarily in Phop-shosite and NetPath). The remaining edges are represented by two directed edges between the proteins involved. Edge weights were assigned by taking the negative logarithm of the associated confidence score, indicating that finding the optimal Steiner Network would be the same as finding the most confident solution (assuming independence between edges). Confidence data was available for the largest of the data sets (InWeb). For HPRD edges that are not in InWeb, we used the minimum nonzero confidence value by default. For the smaller and highly curated data-sets, Phopshosite and NetPath, we used the maximal confidence level.

### Solving DCSN to optimality

#### Definition 6

(Single-Source DCSN) *This is the special case of DCSN in which the demands are precisely* (*a*, *b*_1_, *c*_1_), (*a*, *b*_2_, *c*_2_), …, (*a*, *b_k_*, *c_k_*), *for some* root *a* ∈ *V. We can assume that c*_1_ ≤ *c*_2_ ≤ ⋯ ≤ *c_k_*.

We can derive a natural integer linear program for the Single-Source Directed Condition Steiner Network in terms of network flows, with each demand being met by a flow from source to target:

Each variable *d_uvc_* denotes the flow through edge (*u*,*v*) at condition *c*, if it exists; each variable *d_uv_* denotes whether (*u*,*v*) is ultimately in the chosen solution sub-graph. The first constraint ensures that if an edge is used at any condition, it is chosen as part of the solution. The second constraint enforces flow conservation, and hence that the demands are satisfied, at all nodes and all conditions.

We note that DCSN easily reduces DCSP, as outlined in Theorem 2. However, DCSP is a special case of Single-Source DCSN. Therefore, the Integer Linear Program defined above can be applied to any DCSN instance with a transformation of the instance to DCSP.

### Performance Analysis of Integer Linear Programming

Given the protein protein interaction network *G*, we sample an instance of the node-variant Single-Source DCSN as so[^3^]:

- Instantiate a source node *a*.
- Independently sample *β* nodes reachable from *a*, for each of the *γ* conditions, giving us {*b*_1,1_, …, *b*_*γ*,*β*_}.
- For each node *v* ∈ *V*, include *v* ∈ *V_c_* if *v* lies on the shortest path from *a* to one of {*b*_1,*c*_,.., *b*_*γ*,*c*_}
- For all other nodes *v* ∈ *V_c_* for all *c*, include *v* ∈ *V_c_* with probability *p*.

Using a workstation running an Intel Xeon E5-2690 processor and 250GB of RAM, **optimal solutions** to instances of modest size (generated using the procedure just described) were within reach:

We notice that our primary runtime constraint comes from *γ*, the number of conditions. In practice, the number of conditions does not exceed 100. Therefore, we present a model which can also easily translate and find **optimal solutions** to real world biological problems with practical runtime.

## Conclusion and Discussion

In this paper we introduced the Condition Steiner Network (CSN) problem and its directed variant, in which the goal is to find a minimal subgraph satisfying a set of *k* condition-sensitive connectivity demands. We show, in contrast to known results for traditional Steiner problems, that this problem is NP-hard to approximate to a factor of *C* – *ϵ*, as well as *k* – *ϵ*, for every *C*, *k* ≥ 2 and *ϵ* > 0. We then explored a special case, in which the conditions/graphs satisfy a *monotonicity* property. For such instances we proposed algorithms significantly beating the pessimistic lower bound for the general problem; this was accomplished by reducing the problem to certain traditional Steiner problems. Lastly, we developed and applied an integer programming-based exact algorithm on simulated instances built over the human protein-protein interaction network, and reported feasible runtimes for real-world problem instances.

Importantly, along the way we showed implications of these results for CSN on other network connectivity problems that are commonly used in PPI analysis—such as Shortest Path, Steiner Tree, Prize-Collecting Steiner Tree—when conditions are added. We showed that for each of these problems, we cannot guarantee (in polynomial time) a solution with a value below *C* – *ϵ* times the optimal value. These lower bounds are quite strict, in the sense that naively approximating the problem separately in every condition, and taking the union of those solutions, already gives an approximation ratio of *O*(*C*). At the same time, by relating the various condition Steiner problems to one another, we also obtained some positive results: the condition versions of Shortest Path and Steiner Tree admit good approximations when the conditions are monotonic. Moreover, all of the condition problems (with the exception of Prize-Collecting Steiner Tree) can be solved using a natural integer programming framework that works well in practice.

## Proofs of Main Theorems

### Problem variants

There are several natural ways to formulate the condition Steiner network problem, depending on whether the edges are changing over condition, or the nodes, or both.

#### Definition 10

(Condition Steiner Network (edge variant)) *This is the formulation described in the Introduction: the inputs are G*_1_ = (*V*, *E*_1_), …, *G_C_* = (*V*, *E_C_*), *w*(·), *and* 𝓓 = {(*a_i_*, *b_j_*, *C_j_*)}. *The task is to find a minimum-weight sub-graph 𝓗 C* ⊆ 𝓖 *that satisfies all of the demands*.

#### Definition 11

(Condition Steiner Network (node variant)) *Let the underlying graph be* 𝓖 = (*V*, *E*). *The inputs are G*_1_ = (*V*_1_, *E*(*V_C_*)), …, *G_C_* = (*V_C_*, *E*(*V_C_*)), *w*(·), *and* 𝓓. *Here*, *E*(*V_C_*) ⊆ *E denotes the edges induced by V_C_* ⊆ *V*. *A path satisfies a demand at condition t iff all edges along that path exist in G_c_*.

#### Definition 12

(Condition Steiner Network (node and edge variant)) *The inputs are precisely G*_1_ = (*V*_1_, *E*_1_), …, *G_C_* = (*V_C_*, *E_C_*), *w*(·), *and* 𝓓. *This is the same as the node variant except that each E_c_ can be any subset of E*(*V_c_*).

Similarly, define the corresponding directed problem Directed Condition Steiner Network (DCSN) with the same three variants. The only difference is that the edges are directed, and a demand (*a*, *b*, *c*) must be satisfied by a directed *a* → *b* path in *G_c_*.

The following observation enables all our results to apply to all problem variants.

#### Proposition 2

*The edge*, *node*, *and node-and-edge variants of CSN are mutually polynomial-time reducible via strict reductions (i.e. preserving the approximation ratio exactly). Similarly all three variants of DCSN are mutually strictly reducible.*

#### Proof

The following statements shall hold for both undirected and directed versions. Clearly the node-and-edge variant generalizes the other two. It suffices to show two more directions:

- (Node-and-edge reduces to node) Let (*u*, *v*) be an edge existent at a set of conditions *τ*(*u*, *v*), whose endpoints exist at conditions *τ*(*u*) and *τ*(*v*). To make this a node-condition instance, create an intermediate node *x*_(*u*,*v*)_ existent at conditions *τ*(*u*, *v*), an edge (*u*, *x*_(*u*, *v*)_) with the original weight *w*(*u*, *v*), and an edge (*x*_(*u*, *v*)_, *v*) with zero weight. A solution of cost *W* in the node-and-edge instance corresponds to a node-condition solution of cost *W*, and vice-versa.
- (Node reduces to edge) Let (*u*, *v*) be an edge whose endpoints exist at conditions *τ*(*u*) and *τ*(*v*). To make this an edge-condition instance, let (*u*, *v*) exist at conditions *τ*(*u*, *v*) := *τ*(*u*) ∩ *τ*(*v*). Let every node exist at all conditions; let the edges retain their original weights. A solution of cost *W* in the node-condition instance corresponds to an edge-condition solution of cost *W*, and vice-versa. □

### Inapproximability for general C and k

Here we prove our main theorem, showing optimal hardness for any number of demands. To do this, we introduce a generalization of Label Cover to partite hypergraphs:

#### Definition 13

(*k*-Partite Hypergraph Label Cover (*k*-PHLC)) *An instance of this problem consists of a k-partite*, *k-regular hypergraph G* = (*V*_1_, …, *V_k_*, *E*) *(that is*, *each edge contains exactly one vertex from each of the k parts) and a set of possible labels* Σ. *The input also includes, for each hyperedge e* ∈ *E*, *a* projection function 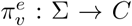 *for each v* ∈ *e*; Π *is the set of all such functions. A labeling of G is a function* 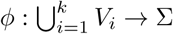 *assigning each node a label. There are two notions of edge satisfaction under a labeling ϕ*:

- *ϕ* strongly satisfies *a hyperedge e* = (*v*_1_, …, *v_k_*) *iff the labels of all its vertices are mapped to the same color*, *i.e.* 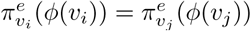 *for all i*, *j* ∈ [*k*].
- *ϕ* weakly satisfies *a hyperedge e* = (*v*_1_, …, *v_k_*) *iff there exists some pair of vertices v_i_*, *v_j_*, *whose labels are mapped to the same color*, *i.e.* 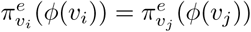 *for some i* ≠ *j* ∈ [*k*].

#### Theorem 7

*For every ϵ* > 0 *and every fixed integer k* ≥ 2, *there is a constant* |Σ| *such that the following promise problem is* **NP**-*hard: Given a k-Partite Hypergraph Label Cover instance* (*G*, Σ, Π), *distinguish between the following cases:*

- *(YES instance) There exists a labeling of G that strongly satisfies every edge.*
- *(NO instance) Every labeling of G weakly satisfies at most ϵ*|*E*| *edges*.

Theorem 7 follows from Feige’s *k*-prover system [34] by taking the number of repetitions to be (a constant depending on *k* and *ϵ*) large enough so that the error probability drops below *ϵ*.

The proof of (*C* – *ϵ*)-hardness and (*k* – *ϵ*)-hardness follows the same outline as the *C* = *k* = 2 case (Theorem 6).

#### Theorem 8

(Main Theorem) *CSN and DCSN are NP-hard to approximate to a factor of C* – *ϵ as well as k* – *ϵ for every fixed k* ≥ 2 *and every constant ϵ* > 0. *For DCSN*, *this holds even when the underlying graph is acyclic.*

#### Proof

Given the *k*-PHLC instance in the form (*G* = (*V*_1_, …, *V_k_*, *E*), Σ, Π), and letting *v*_*c*,*i*_ denote the ith node in *V_c_*, construct a DCSN instance (𝓖 = (*G*_1_, …, *G_k_*), along with *k* demands) as follows. For every *c* ∈ [*k*], create nodes 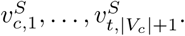 Create a *v*_*c*,*i*_-*bundle* from each 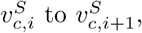 whose *ℓ*-strands (for *ℓ* ∈ Σ) are each a chain of bundles, one for each incident hyperedge *e* = (*v*_1, *i*_1__, …, *v*_*c*, *i*_, …, *v*_*k*, *i*_*k*__) ∈ *E*. Each (*v*_1, *i*_1__, …, *v*_*c*,*i*_, …, *v*_*k*, *i*_*k*__)-bundle has a (*v*_1, *i*_1__,… *ℓ*_1_, …, *v*_*c*,*i*_,*ℓ_c_*, …, *v*_*k*,*i*_*k*__,*ℓ_k_*)-path for each agreeing combination of labels—that is, every *k*-tuple (*ℓ*_1_, …, *ℓ_c_*, …, *ℓ_k_*) such that: 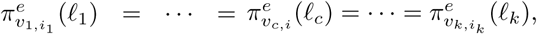 where *e* is the shared edge. If there are no such combinations, then the *e*-bundle is a single simple strand.

For *c* ∈ [*k*], set all the edges in the *v*_*c*,*i*_-bundles to exist in *G_c_* only. Now, for each (*v*_1,*i*_l__, *ℓ*_1_, …, *v*_*k*,*i*_*k*__,*ℓ_k_*), merge together the (*v*_1,*i*_l__, *ℓ*_1_, …, *v*_*k*,*i*_*k*__, *ℓ_k_*)-paths across all G_c_ that have such a strand. Finally, the connectivity demands are 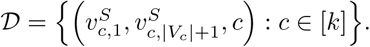

The analysis follows the *k* = 2 case. Suppose we have a YES instance of *k*-PHLC, with optimal labeling 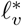 to each node 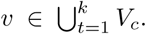 Then an optimal solution 𝓗^∗^ to the constructed DCSN instance is to traverse, at each condition *c* and for each *v*_*c*,*i*_-bundle, the path through the 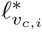-strand. In particular for each (*v*_1,*i*_l__, …, *v*_*k*, *i*_*k*__)-bundle in that strand, traverse the 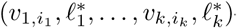-path.

In tallying the total edge cost, 𝓗^∗^ ∩ *G*_1_ (the subgraph at condition 1) incurs a cost of |*E*|, one for each contact edge. The sub-graphs of 𝓗^∗^ at conditions 2, …, *k* account for no additional cost, since all contact edges correspond to a label which agrees with all its neighbors’ labels, and hence were merged with the agreeing contact edges in the other sub-graphs.

Conversely suppose we have a NO instance of *k*-PHLC, so that for any labeling 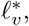 for at least (1 – *ϵ*)|*E*| hyperedges *e*, the projection functions of all nodes in *e* disagree. By definition, any solution to the constructed DCSN instance contains a simple 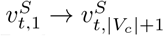 *P_c_* at each condition *c*. As before, *P*_1_ alone incurs a cost of exactly |*E*|. However, at least (1 – *ϵ*)|*E*| of the hyperedges in *G* cannot be weakly satisfied; for these hyperedges *e*, for every pair of neighbors *v*_*c*,*i*_*c*__, *v*_*c*′,*i*_*c*′__, ∈ *e*, there is no path through the *e*-bundle in 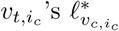 - strand that is merged with any of the paths through the *e*-bundle in 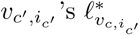-strand (for otherwise, it would indicate a labeling that weakly satisfies *e* in the *k*-PHLC instance). Therefore paths *P*_2_, …, *P_k_* each contribute at least (1 – *ϵ*)|*E*| additional cost, so the solution has total cost at least (1 – *ϵ*)|*E*| · *k*.

It follows from the gap between the YES and NO cases that DCSN is NP-hard to approximate to within a factor of *k* – *ϵ* for every constant *ϵ* > 0; and since *C* = *k* in our construction, it is also NP-hard for *C* – *ϵ*. Moreover since The directed condition graph we constructed is acyclic, this result holds even on DAGs. As before, the same analysis holds for the undirected problem CSN by undirecting the edges. □

### Explicit algorithm for Monotonic Single-Source DCSN

We provide a modified version of the approximation algorithm presented in Charikar et al. [21] for Directed Steiner Tree (DST), which achieves the same approximation ratio for our problem Monotonic Single-Source DCSN.

We provide a similar explanation as of that presented in Charikar et al. Consider a trivial approximation algorithm, where we take the shortest path from the source to each individual target. Consider the example where there are edges of cost *C* – *ϵ* to each target, and a vertex *v* with distance *C* from the source, and with distance 0 to each target. In such a case, this trivial approximation algorithm will achieve only an Ω(*k*)-approximation. Consider instead an algorithm which found, from the root, an intermediary vertex *v*, which was connected to all the targets via shortest path. In the case of the above example, this would find us the optimal sub-graph. The algorithm below generalizes this process, by progressively finding optimal substructures with good cost relative to the number of targets connected. We show that this algorithm provides a good approximation ratio.

#### Definition 14

(Metric closure of a condition graph) *For a directed condition graph* 𝓖 = (*G*1 = (*V*, *E*_1_), *G*_2_ = (*V*, *E*_2_), …, *G_C_* = (*V*, *E_C_*)), *define its metric closure to be G* = (*V*, *E*, *w̃*) *where E* = ⋃_*c*_ *E_c_ and w̃*(*u*, *v*, *c*) *is the length of the shortest u* → *v path in G_c_ (note that in contrast with w*, *w̃ takes three arguments)*.

#### Definition 15

(*V*(*T*)) *Let T be a tree with root *r*. We say a demand of the form* (*r*, *b*, *c*) *is satisfied by T if there is a path in T from r to b at condition c. V* (*T*) *is then the set of demands satisfied by T*.

#### Definition 16

(*D*(*T*)) *The density of a tree T is* 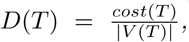 *where cost*(*T*) *is the sum of edge weights of T*.

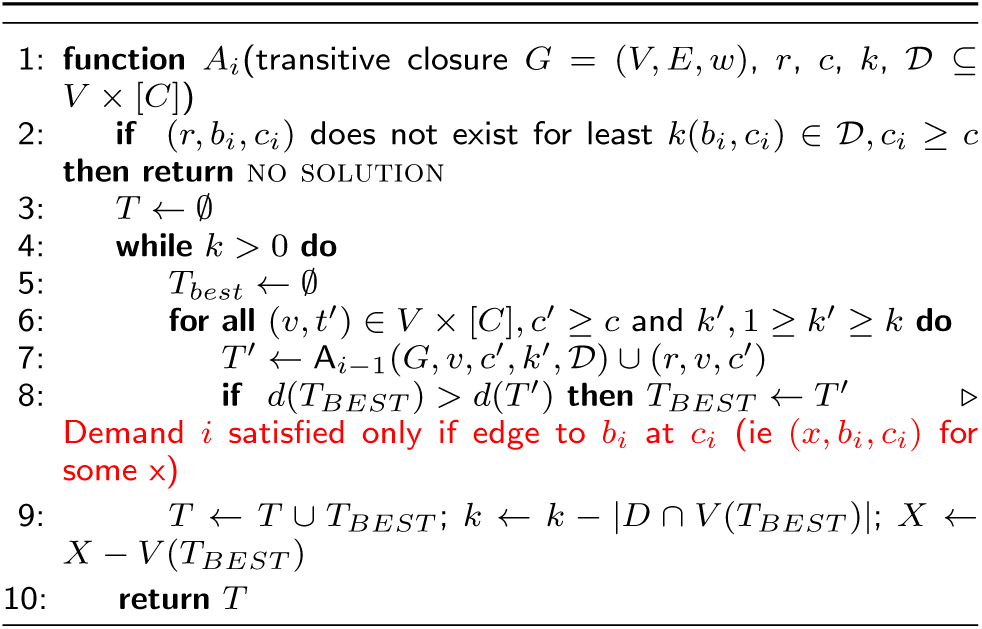

The way we will prove the approximation ratio of this algorithm is to show that it behaves precisely as the algorithm of Charikar et al. does, when given as input the DST instance produced by our reduction from Monotonic Single Source DCSN (Lemma 3).

#### Proposition 3

*The algorithm above is equivalent to the algorithm of Charikar et al., when applied to the DST instance output by the reduction of Lemma 3*.

#### Proof

To see this, note that in our reduced instance, we see a collection of vertices, *v*^1^, …, *v*^|*C*|^ Therefore, the only equivalent modifications needed to the original algorithm are:

- In the input, rather than keeping track of the current root as some vertex *v^i^*, keep track of *v* at the current condition instead, i.e. (*v*,*i*).
- The distance from some *v^i^* to *x^j^*, *j* ≥ *i* is simply the distance from *v* to *x* at condition *j*, i.e. *w̃*(*v*,*x*,*j*).
- Instead of looping through all vertices in the form *v*^1^, …, *v*^|*C*|^, we instead loop through all vertices, and all conditions.

Therefore this algorithm guarantees the same approximation ratio for Monotonic Single Source DCSN as the original algorithm achieved for DST. In particular for all *i* > 1, *A_i_*(*G*, *a*,0, *k*, *D*) provides an *i*^2^(*i* – 1)*k*^1/*i*^ approximation to DCSN, in time *O*(*n^i^k*^2*i*^) [21, 35][^4^]. □

## List of Abbreviations

CPCST: Condition Prize-Collecting Steiner Tree
CSN: Condition Steiner Network
CST: Condition Steiner Tree
CSP: Condition Shortest Path
DSN: Directed Steiner Network
DST: Directed Steiner Tree
DCSN: Directed Condition Steiner Network
DCSP: Directed Condition Shortest Path
*k*-PHLC: *k*-Partite Hypergraph Label Cover
MKL: Minimum k-Labeling
PPI: Protein Protein Interaction

## Declarations

### Ethics approval and consent to participate

Not applicable

### Consent for publication

Not applicable

### Availability of data and material

Our simulation code can be found at the following URL: https://github.com/YosefLab/condition_connectivity_problems

## Author’s contributions

All authors conceived and designed the study. JW and BW derived the main hardness results. AK derived the monotonic hardness results and approximation algorithm. NY was the PI and oversaw the project.

## Competing interests

The authors declare that they have no competing interests.

## Funding and Acknowledgements

This work was partially supported by the National Science Foundation Graduate Research Fellowship Program award DGE 1106400, NIH grants U01HG007910 and U01MH105979, and the U.S.-Israel Binational Science Foundation.

## Footnotes

1 *V* is the set of nodes in the reference graph *G*

2 Throughout this paper, *n* := |*V*| denotes the number of nodes in the relevant graph.

4 As previously mentioned, this variant reduces to the edge variant via reduction, and vice versa

4 The first paper [21] incorrectly claims a bound of *i*(*i* – 1)*k*^1/*i*^; this was corrected in [35].

